# Evaluation of gelatin bloom strength on gelatin methacryloyl hydrogel properties

**DOI:** 10.1101/2023.11.13.566924

**Authors:** Samantha G. Zambuto, Samyuktha S. Kolluru, Eya Ferchichi, Hannah F. Rudewick, Daniella M. Fodera, Kristin M. Myers, Silviya P. Zustiak, Michelle L. Oyen

**Affiliations:** Department of Obstetrics and Gynecology, Washington University School of Medicine, St. Louis, MO 63130; Department of Biomedical Engineering, Washington University in St. Louis, St. Louis, MO 63130; Center for Women’s Health Engineering, Washington University in St. Louis, St. Louis, MO 63130; The Institute of Materials Science & Engineering, Washington University in St. Louis, St. Louis, MO 63130; Department of Biomedical Engineering, Saint Louis University, St. Louis, MO 63103; Department of Chemical Engineering, Texas A&M University, College Station, TX 77843; Department of Biomedical Engineering, Columbia University, New York, NY; Department of Mechanical Engineering, Columbia University, New York, NY

**Keywords:** gelatin, gelatin methacryloyl, bloom strength, hydrogel properties, nanoindentation, trophoblast spheroids

## Abstract

Gelatin methacryloyl (GelMA) hydrogels are widely used for a variety of tissue engineering applications. The properties of gelatin can affect the mechanical properties of gelatin gels; however, the role of gelatin properties such as bloom strength on GelMA hydrogels has not yet been explored. Bloom strength is a food industry standard for describing the quality of gelatin, where higher bloom strength is associated with higher gelatin molecular weight. Here, we evaluate the role of bloom strength on GelMA hydrogel mechanical properties. We determined that both bloom strength of gelatin and weight percent of GelMA influenced both stiffness and viscoelastic ratio; however, only bloom strength affected diffusivity, permeability, and pore size. With this library of GelMA hydrogels of varying properties, we then encapsulated Swan71 trophoblast spheroids in these hydrogel variants to assess how bloom strength affects trophoblast spheroid morphology. Overall, we observed a decreasing trend of spheroid area and Feret diameter as bloom strength increased. In identifying clear relationships between bloom strength, hydrogel mechanical properties, and trophoblast spheroid morphology, we demonstrate that bloom strength should considered when designing tissue engineered constructs.

## 1. Introduction

Gelatin methacryloyl (GelMA) hydrogels are used in biomaterials research for a plethora of biomedical applications, spanning from basic science studies to tissue engineering, regenerative medicine, bioprinting, and drug delivery. GelMA is readily commercially available and exhibits exceptional consistency^1,2^. GelMA is prepared from gelatin, a soluble polypeptide mixture of partially hydrolyzed collagens derived from a variety of tissue sources, including skin, tendons, and bones^3^. Gelatin is sourced primarily from porcine, bovine, and piscine tissues^3^, but certain vendors now carry human-derived gelatins as well. Gelatin, as well as GelMA, is a naturally derived polymer that consists primarily of collagen types I and III, with trace amounts of other collagen sub-types^3^. GelMA also retains the arginine-glycine-aspartic acid (RGD) cell binding motifs and MMP-sensitive degradation sites in gelatin^2,4^, rendering this material bioactive, biocompatible, and biomimetic.

Gelatin hydrogels are traditionally polymerized via physical crosslinking and form thermo-reversible hydrogels via physical interactions between collagen molecules^3^. Thus, gelatin hydrogels are unstable at physiological temperatures and can be heterogeneous in terms of their mechanical properties^3^. To ameliorate these limitations, gelatin can be chemically functionalized as GelMA which allows for photopolymerization and greater tunability and control over hydrogel mechanical properties^3^. Hydrogel mechanical properties can be tuned by varying the degree of functionalization, polymer weight percent, photo-initiator concentration, and UV crosslinking time^3^. Previous studies have suggested that the hydrogel composition can influence cell behavior and hydrogel characteristics may need to be optimized to elicit physiologically relevant cell behavior^3,5^.

The effect of gelatin bloom strength (Blst) on GelMA hydrogel properties has not yet been explored. Unlike other polymers which are differentiated by their molecular weight, gelatin is differentiated by bloom strength because the hydrolytic conversion of collagen produces a distribution of gelatin molecular weights from less than 10,000 g/mol to over 400,000 g/mol^6,7^. The Bloom number corresponds to average molecular mass, where low bloom corresponds to lower (20 – 25 kDa) molecular mass and high bloom corresponds to higher (50 – 100 kDa) molecular mass. Bloom strength of gelatin is quantified by dropping a standardized 0.5 inch diameter plunger a pre-specified distance of 4 mm into a gelatin gel and determining the force required impress the gel in grams^7–9^. The weight in grams from the machine corresponds to bloom strength of the gel, where a higher force results in a higher bloom strength^7,9^. Bloom strength typically ranges from 50 to 300 g and is dependent on many factors, including gelatin structure and molecular mass^6^. Bloom strength (*B*) and solution concentration (*C*) both affect the properties of gelatin and hence^6^, these two factors were combined into a single property known as gel strength (*GS*): *GS* = *C*^2^*B*. In the food industry, two gelatin batches of equal gel strength are considered interchangeable^6^. Higher bloom strength gelatin is considered higher quality than lower strength gelatin because it has longer collagen chains and contains fewer impurities^8^. High bloom strength also corresponds to higher melting and gelling temperatures which results in shorter gel time. Gel strength has been shown to linearly correlate with the triple-helical content and the compressive modulus of gelatin^6,8^; however, the role of gelatin bloom strength on GelMA mechanical properties has not yet been defined.

Our goal was to define the role of bloom strength on GelMA mechanical properties and subsequently investigate the role of these properties on behavior of encapsulated cells. For these studies, we used the Swan 71 immortalized trophoblast cell line because of its relevance to understanding placental function. With these GelMA hydrogel variants, we created and categorized a library of hydrogels so we could methodically define how bloom strength influenced trophoblast spheroid morphology. We synthesized three GelMA variants of varying bloom strength (Low, Medium, High) and systematically characterized hydrogel mechanical properties through spherical nanoindentation. We then encapsulated trophoblast spheroids into these hydrogel variants and quantified spheroid area, Feret diameter, and circularity. We identified clear relationships between bloom strength, hydrogel mechanical properties, and trophoblast spheroid morphology. Here, we highlight the bloom strength effects on GelMA hydrogel properties and cellular behavior.

## 2. Materials and Methods

### 2.1. Gelatin methacryloyl (GelMA) synthesis and characterization

#### 2.1.1. GelMA Synthesis

Gelatin methacryloyl (GelMA) was synthesized using the one pot method developed by Shirahama et al ^1^. Briefly, gelatin type A (acid-hydrolyzed; porcine-derived) of varying Blst (“Low” 80-120 g, Sigma Aldrich G6144; “Medium” ∼175 g, Sigma Aldrich G2625; and “High” ∼300 g Sigma Aldrich G2500) were dissolved 10% w/v in carbonate buffer (ThermoFisher) 28382 at 50°C. Methacrylic anhydride (40 *μ* L/g gelatin; Sigma Aldrich 276685) was added dropwise to the solution while stirring and the reaction was allowed to proceed for 1h. The reaction was subsequently stopped with the addition of deionized water (40 mL/g gelatin) and the pH of the solution was adjusted to 6 - 7. The solution was transferred to dialysis membranes (12 -14 kDa molecular weight cut off; Fisher Scientific 21-152-8) and dialyzed against deionized water for 7 days and subsequently lyophilized (LABCONCO).

#### 2.1.2. Environmental Scanning Electron Micrograph Imaging

Environmental Scanning electron micrograph (ESEM) images were taken for GelMA variants. Lyophilized GelMA samples were attached to a pinstub via carbon tape and imaged using an Environmental Scanning Electron Microscope Thermofisher Quattro S ESEM. Instrument settings are as follows: 3.0 Spot Size, 250X magnification, and 2.00 kV voltage.

#### 2.1.3. Nuclear Magnetic Resonance Spectroscopy

The lyophilized GelMA was dissolved in Deuterium water (20 mg/mL) then measured by proton nuclear magnetic resonance (^1^H-NMR) spectroscopy (Bruker Avance III HD 700 MHz). Degree of functionalization (DOF_NMR_) was calculated by comparing the lysine methylation (ME) peaks (2.9 ppm) of GelMA to gelatin using **Eq. 1**.

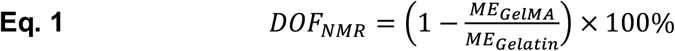

#### 2.1.4. Attenuated Total Reflectance-Fourier Transform Infrared Spectroscopy

Lyophilized GelMA samples and powdered stock gelatin samples were analyzed via attenuated total reflectance-Fourier transform infrared spectroscopy (ATR-FTIR) using a Bruker Alpha II FTIR outfitted with a zinc selenide crystal (spectral range 20,000 – 500 cm^-1^). Spectra were averaged over 24 scans and were recorded from 600 – 4000 cm^-1^ with a resolution of 4 cm^-1^. A background scan was performed prior to each sample measurement. Measurements were repeated three times with a five second gap between each measurement. The three scans were then averaged and normalized in Prism 9 software. Raw and normalized, non-truncated data can be found in **Supplemental Figure 1**.

### 2.2. GelMA hydrogel fabrication and characterization

#### 2.2.1. Hydrogel Fabrication

The 5 wt% (weight percent) and 10 wt% hydrogels were prepared by dissolving GelMA in phosphate buffered saline (PBS) at 37°C. Once the solution was dissolved, 0.1% w/v lithium acylphosphinate (LAP; Sigma Aldrich 900889-1G) photoinitiator was added to the solution and thoroughly mixed. Prepolymer solution was pipetted (20 *μ* L or 100 *μ* L) into custom Teflon molds (5 mm diam., 1 mm height or 10 mm diam., 1 mm height, respectively). Hydrogels were polymerized for 5 min under UV light (λ = 365 nm; Spectroline EA-160) and subsequently removed from the molds. All experiments used 5 mm diam. gels except for mass swelling experiments for which the 10 mm diam. gels were used. Despite different diameters, the heights of the gels were kept consistent to ensure the same UV penetration.

#### 2.2.2. ATR-FTIR

GelMA hydrogels were synthesized and analyzed via ATR-FTIR as described above. Hydrated (overnight in PBS) and lyophilized hydrogels were then analyzed via ATR-FTIR. Raw versus normalized data (full spectrum and truncated) can be found in **Supplemental Figures 2 and 3**.

#### 2.2.3. Nanoindentation

Spherical nanoindentation (Piuma, Optics11Life) was used to quantify the time-dependent material properties of GelMA hydrogel variants. A 26 *μ* m probe radius (*R*) with a cantilever stiffness of 0.42 N/m stiffness was used for indentation. In preparation for testing, GelMA hydrogels were cast on GelBond® PAG Film (Lonza 54746) and the GelBond® was superglued (The Original SuperGlue Corporation) to glass Petri dishes (Pyrex). Hydrogels were swelled overnight at room temperature in 1X PBS or Opti-Free Contact Lens Solution. Prior to testing, samples were submerged in Opti-Free Contact Lens Solution and were allowed to re-equilibrate for at least 20 min prior to testing. Opti-Free solution was used because it contains surfactants which reduce adhesion between the probe and hydrogel^10^. Hydrogels were indented to a fixed depth of 3 *μ* m (*δ*_0_) under displacement control. After a 2 s ramp time to the prescribed indentation depth, the probe position was held for 15 s to yield a load-relaxation curve that approached equilibrium. At minimum, three hydrogels were tested per condition. A 5x5 matrix scan (25 indentation points) with 250 *μ* m between each indentation point was performed, yielding 55 – 139 individual indentation points per condition from 5 – 25 points per individual hydrogel sample.

#### 2.2.4. Nanoindentation Data Analysis

Time-dependent hydrogel properties were determined by fitting load-relaxation curves with an established poroelastic-viscoelastic (*P*_*PVE*_) model in MATLAB as previously described^11–16^. Load-relaxation curves were fit with the *PVE* model to determine stiffness (*E*), viscoelastic ratio (*G*_∞_/*G*_0_), intrinsic permeability (*k*), and diffusivity (*D*). Fitted data were excluded from the final data set if the load-relaxation curve displayed sharp discontinuities, increasing loads over time, or Δ*P* < 0.005.

The combined poroelastic-viscoelastic effect (*P*_*PVE*_) is described by the poroelastic (*P*_*PE*_), viscoelastic (*P*_*VE*_), and elastic load (*P*_∞_) responses calculated from equilibrium properties^16^:

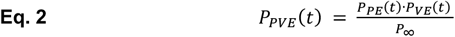

The poroelastic response (*P*_*PE*_) is derived as:

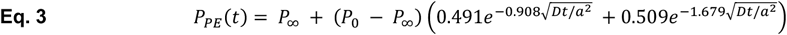

Where *D* is the diffusion coefficient, *a* is the contact radius derived from the probe radius (*R*) and applied indentation depth (δ_0_) by *a*^2^ = *R* ·δ_0_.

Intrinsic permeability *k* was then calculated as:

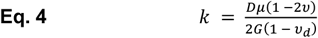

where *μ* is the solvent viscosity (*μ* = 0.89 × 10^−3^ *Pa* · *s*), *v* is Poisson’s ratio set as *v* = 0.5 which is a commonly assumed value for macro-scale hydrated testing of hydrogels where the solvent migration is negligible during ramp loading, is *v*_*d*_ drained Poisson’s ratio, and *G* is the shear modulus. Pore size (ξ) was inferred from permeability by 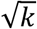.

The viscoelastic response (*P*_*VE*_) is calculated using a generalized Maxwell model^11^, consisting of a linear spring connected in parallel with *n* number of Maxwell units (serial connection of a linear spring and dashpot; n = 2 in this case):

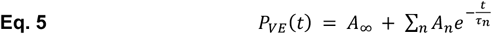

Where *A*_∞_ and *A*_*n*_ are empirical constants and *τ*_*n*_ is the characteristic relaxation time of the *n*^th^ Maxwell element. The coefficients can approximately be related as follows:

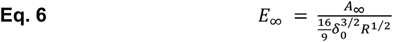

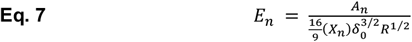

Where *E*_∞_ is the equilibrium elastic modulus, *E*_*n*_ is the elastic modulus, *X*_*n*_ is the ramp correction factor which accounts for finite ramp time (*t*_*r*_) and is defined as:

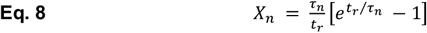

The viscoelastic component of the model is thus defined as:

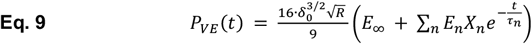

From these equations, we can calculate the stiffness, *E*, as: *E* = 3*G* and diffusivity, *D*, as: 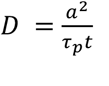. Best fit lines relating Blst to stiffness and viscoelastic ratio were created using the non-linear fit function in Prism 9 set to a lognormal model.

#### 2.2.5. Mass Swelling Ratio

Mass swelling ratio was determined by comparing the swollen (*m*_*s*_) and dry (*m*_*d*_) hydrogel mass using **Eq. 10**. Hydrogels were swollen to equilibrium, weighed, lyophilized, and weighed once more to determined dry mass.

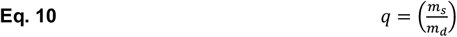

#### 2.2.5. Swan71 Spheroid Encapsulation and Analysis

Swan71 trophoblast cells (Applied Biological Materials Inc. T0532) were cultured on T25 flasks in phenol red-free DMEM/F12 (Gibco 21041-025) supplemented with 10% charcoal-stripped fetal bovine serum, 1% penicillin/streptomycin (Gibco 15140-122), and 500 ng/mL puromycin (Sigma Aldrich, P8833). Once the cells reached 80-90% confluence, 4,000 cells were added to each well of a 96 well round bottom plate (200 *μ* L/well; Corning 4515) and left undisturbed for 24 h. After 24 h, individual spheroids were encapsulated in GelMA hydrogels as previously described^17–19^. Spheroid-laden hydrogels were cultured in 48 well plates with 800 *μ* L of medium containing a reduced amount of fetal bovine serum (2%). Spheroids were imaged following encapsulation in Bright Field on a Keyence All-in-One Fluorescence Microscope BZ-X series with a 10X objective. All cell cultures were grown in 5% CO_2_ incubators at 37°C.

Spheroid images were segmented using FIJI^20^. A FIJI macro was developed to quantify spheroid area, Feret diameter, and circularity as follows (**Supplemental Figure 4**): images were converted into 8-bit and then binarized, the edges of the spheroid were identified and dilated to reduce the roughness of the outline, any holes within the binarized shape were filled using erosion to revert the spheroid back to its original size, the scale for the image was set, and then the command “Analyze Particles” was used to obtain the area, Feret diameter, and circularity. A best fit line relating Blst to spheroid area and Feret diameter were created using the Lognormal Non-Linear Fit option in Prism 9.

### 2.3. Statistical Analysis

Nanoindentation data (stiffness, viscoelastic ratio, permeability, diffusivity, pore size), mass swelling ratio, and trophoblast spheroid data were analyzed using ordinary two-way ANOVAs assuming normal data distribution and homoscedasticity and Tukey’s multiple comparisons post-hoc tests using Prism 9 software. Because of these assumptions, we set a more conservative significance value of *p* < 0.01. All plots except the ^1^H-NMR plots were created in Prism 9 software.

## 3. Results

### 3.1. Synthesis of GelMA Blst Variants

We synthesized three variants of GelMA (Low Blst, Medium Blst, and High Blst) and characterized their morphology, chemical properties, and composition (**Fig. 1**). From ESEM images, GelMA Medium Blst and High Blst had similar morphology whereas GelMA Low Blst had more strand-like regions compared to the other two variants (**Fig. 2A**). Next, we sought to verify the synthesis reaction and determine degree of functionalization. From ^1^H-NMR, we observed the addition of new proton peaks between 6 - 5 ppm corresponding to methacrylation of gelatin and reduction in the 2.8 ppm peak from gelatin to GelMA which corresponds to a reduction of the free lysine groups and successful modification (**Fig. 2B**)^1–3^. From the ^1^H-NMR spectra, we quantified the degree of functionalization of the gelatin backbone for each variant (**Table 1**), and determined each variant was within a similar range for degree of functionalization. We then performed compositional analysis via ATR-FTIR for gelatin and GelMA. We observed characteristic gelatin peaks around 1600 cm^-1^ associated with amide I C=O stretching, 1500 cm^-1^ associated with N-H bending coupled to C-H stretching, and 1200 cm^-1^ associated with amide III C-N stretching and N-H bending^21^. Compositional analysis via ATR-FTIR demonstrated differences between GelMA and gelatin corresponding to the amide I peak (∼1600 nm), amide II peak (∼1500 nm), and amide III peak (∼1200 nm) which indicate successful methacrylation of gelatin (**Fig. 2C**).

**Table 1.** Degree of modification of gelatin methacryloyl variants calculated from^1^H-NMR.

| GelMA Bloom Strength | $^1\text{H-NMR}$ |
| --- | --- |
| Low | 30.8% |
| Medium | 44.2% |
| High | 42.9% |

**Figure 1.**
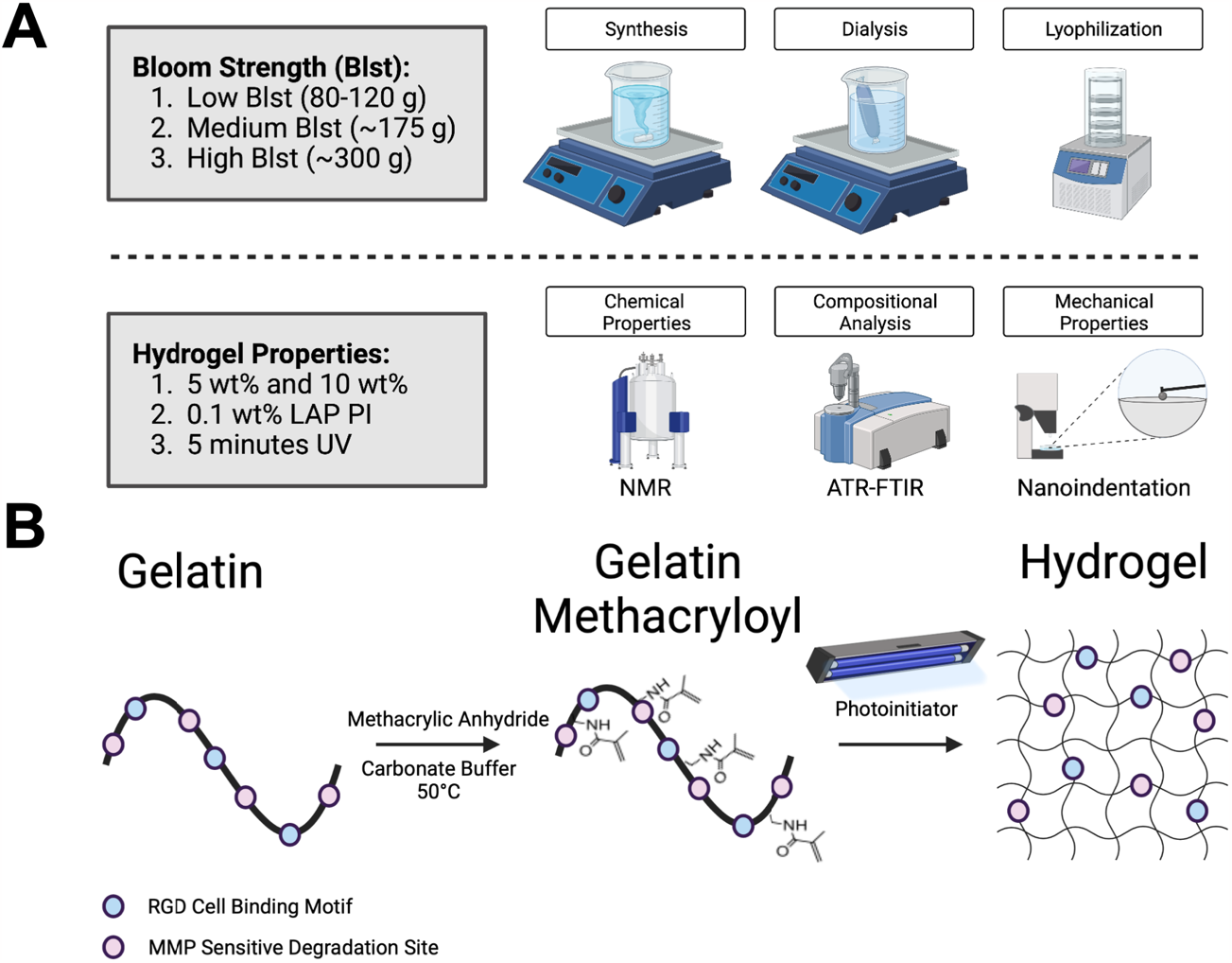
Experimental schematic of **A**. Gelatin methacryloyl (GelMA) synthesis and GelMA hydrogel characterization. **B**. Chemical synthesis and hydrogel fabrication reaction. Created with BioRender.com
.

**Figure 2.**
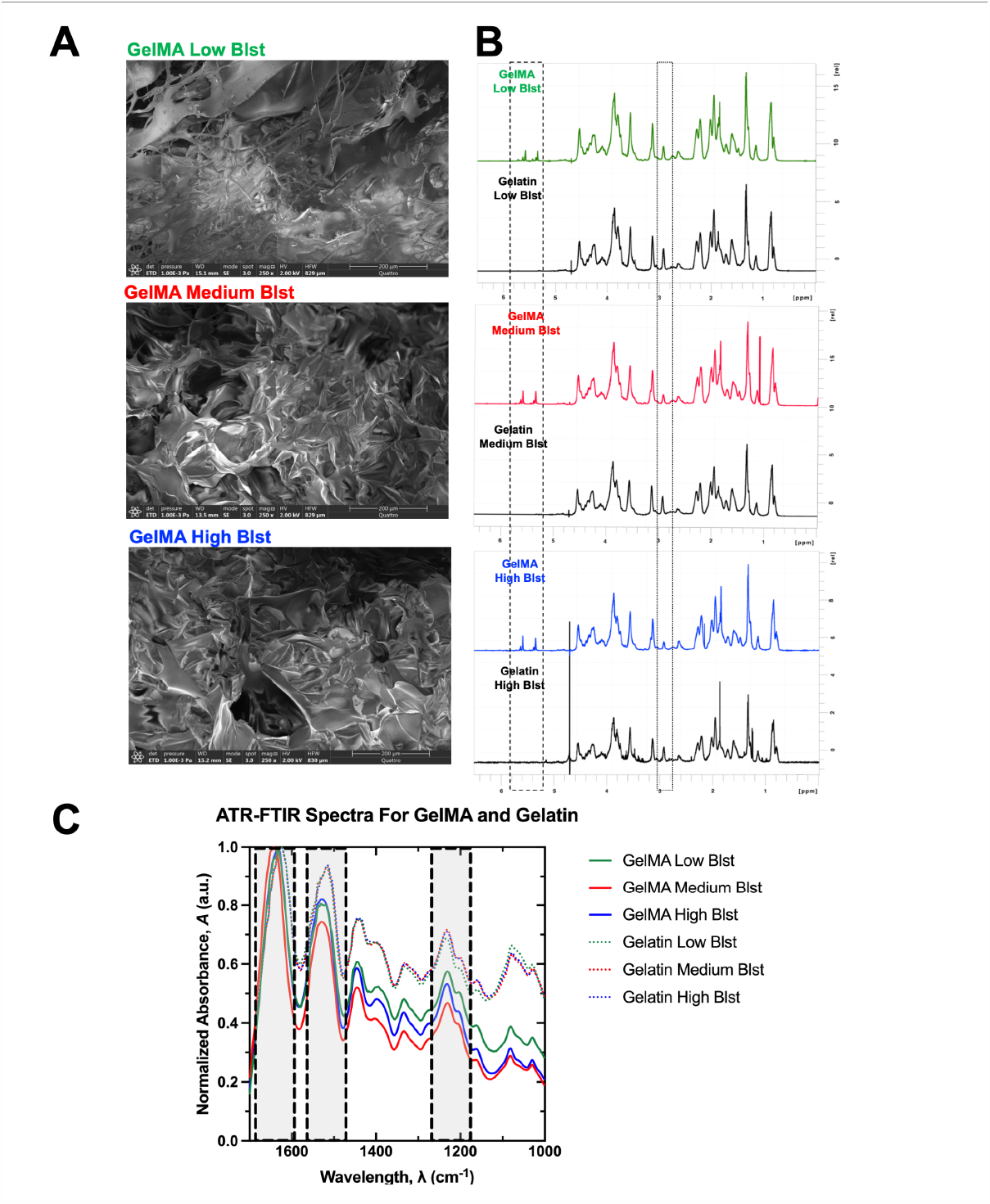
**A**. ESEM Images of GelMA variants. Scale Bar: 200 *μ* m. **B**. ^**1**^H-NMR Spectra for GelMA variants. Chemical shifts confirming presence of acrylic protons of the methacrylate group between 5.35 and 5.65 ppm and reduction of lysine methylene peak at 2.8 ppm. **C**. Truncated (1000-1700 cm-1) normalized ATR-FTIR spectra for gelatin and GelMA variants. Shaded regions highlight peak differences between GelMA and gelatin.

### 3.2. Time-Dependent Properties of GelMA Blst Hydrogel Variants

We next fabricated 5 wt% and 10 wt% GelMA hydrogels for each Blst, thus creating a library of hydrogels with varying GelMA Blst, concentration, and gel strength. The 5 wt% Low Blst hydrogel variants did not polymerize even after 10 minutes of UV exposure and therefore, could not be mechanically characterized. Using spherical nanoindentation, we characterized the time-dependent properties of GelMA Blst hydrogel variants (**Fig. 3**; n = 3 – 6 hydrogels; n = 55 – 139 individual indentation points for each condition). Blst (*p* < 0.0001) and weight percent (*p* < 0.0001) statistically significantly affected stiffness (**Fig. 3A**). We observed an increasing trend in stiffness as Blst and weight percent increased (**Table 2**). Viscoelastic ratio followed a similar trend, for which Blst (*p* < 0.0001) and weight percent (*p* < 0.0001) statistically significantly affected viscoelastic ratio and the ratio increased with Blst and weight percent (**Fig. 3B**).Diffusivity (**Fig. 3C**) and permeability (**Fig. 3D**) also followed similar trends for which Blst significantly affected permeability (*p* < 0.0001) but weight percent did not (*p* = 0.7703). Blst significantly affected diffusivity (*p* = 0.0013) but weight percent did not (*p* = 0.0292). The Low Blst variant was statistically significantly different compared to all groups for permeability and all groups except High Blst 10 wt% for diffusivity. The data were also more variable across a wider range for the Low Blst hydrogel variant compared to Medium and High Blst hydrogels.

**Table 2.** Gel strength of gelatin methacryloyl hydrogel variants calculated from the equation *GS* = *C*^2^*B*.

| GelMA Bloom Strength | Bloom, B (g) | Concentration, C (wt%) | Gel Strength, GS=C<sup>2</sup>B (unitless) | Measured Stiffness E (kPa) |
| --- | --- | --- | --- | --- |
| Low | 100 | 10 | 10,000 | 0.1132 |
| Medium | 175 | 5 | 4,375 | 0.1751 |
| Medium | 175 | 10 | 17,500 | 0.1896 |
| High | 300 | 5 | 7,500 | 0.1575 |
| High | 300 | 10 | 30,000 | 0.2093 |

**Figure 3.**
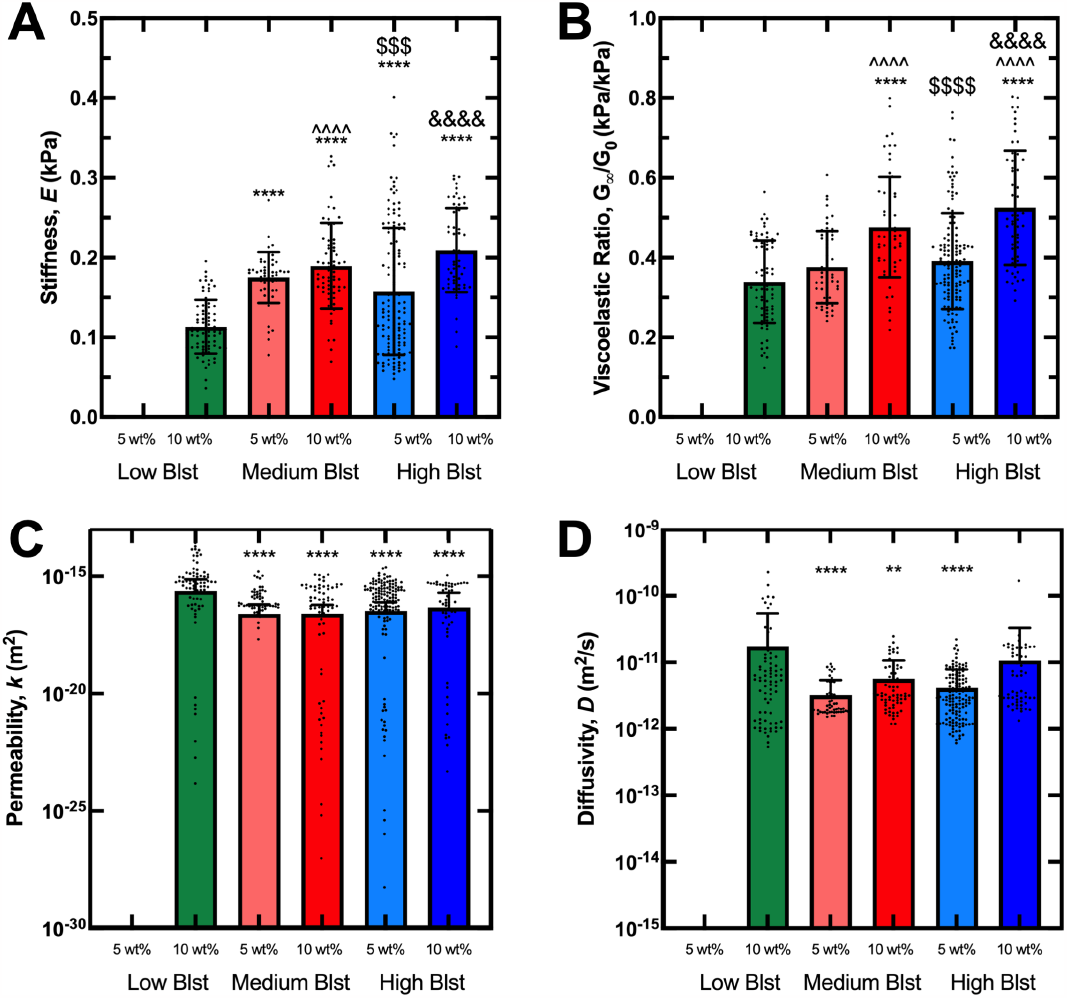
Time-dependent hydrogel properties determined by fitting load-relaxation curves obtained via spherical nanoindentation with a combined poroelastic-viscoelastic model. **A**. Stiffness, **B**. Viscoelastic ratio, **C**. Permeability, and**D**. Diffusivity. Data depicted as mean ± standard deviation for stiffness and viscoelastic ratio and mean + standard deviation for diffusivity and permeability. Each plot depicts individual nanoindentation points compiled from hydrogel samples (n = 55 – 139 individual points per condition). Data analyzed using an ordinary main effects only two-way ANOVA and Tukey posthoc with significance set to p < 0.01. *: different from low blst 10 wt%, ** p < 0.01, **** p < 0.0001 ^: different from med blst 5 wt%, ^^^^ p < 0.0001 $:different from med blst 10 wt%, $$$ p < 0.001, $$$$ p < 0.0001 &: different from high blst 5 wt%, &&&& p < 0.0001

### 3.3. Crosslinking Density and Pore Size of GelMA Blst Hydrogel Variants

We next sought to assess hydrogel crosslinking density as well as hydrogel pore size. Mass swelling ratio (**Fig. 4A**), an indication of degree of crosslinking, decreased with increasing polymer weight percent for each Blst. Blst (Two-way ANOVA, *p* = 0.41) had no effect on crosslinking density; however, weight percent (Two-way ANOVA, *p* < 0.0001) significantly affected hydrogel crosslinking density. We inferred pore size from permeability nanoindentation data (**Fig. 4B**). Blst significantly affected pore size (Two-way ANOVA, *p* < 0.0001) but weight percent did not. The Low Blst hydrogel variant was significantly different from all groups (*p* < 0.0001 all groups). Pore size was approximately 4 nm across the Medium and High Blst conditions, regardless of weight percent. The pore size of Low Blst hydrogels was highly variable, with a maximum value of 52.6 nm and an average value of 10.9 nm.

**Figure 4.**
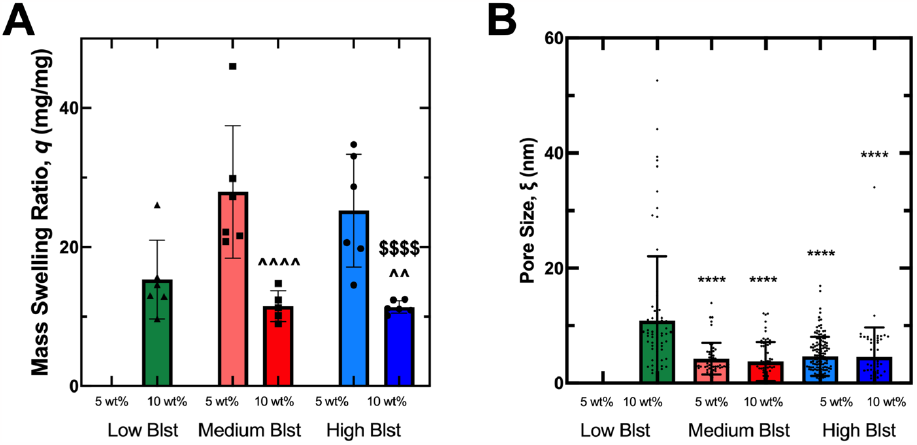
Crosslinking density and pore size for GelMA hydrogel variants. **A**. Mass swelling ratio (n=5-6 hydrogels per condition). **B**. Pore size calculated from permeability nanoindentation data. Plot depicts individual pore size values from nanoindentation points compiled from hydrogel samples (n = 55 – 139 individual points per condition). Data presented as mean ± standard deviation. Data analyzed using an ordinary main effects only two-way ANOVA and Tukey posthoc with significance set to p < 0.01 *: different from low blst 10 wt%, **** p < 0.0001 ^: different from med blst 5 wt%, ^^ p < 0.01, ^^^^ p < 0.0001 $: different from med blst 10 wt%, $$$$ p < 0.0001

### 3.4. Quantification of Trophoblast Spheroid Morphology in GelMA Blst Hydrogel Variants

Next, we sought to determine if Blst affected cellular properties. We seeded single Swan71 trophoblast spheroids per hydrogel and imaged the spheroids immediately after encapsulation (**Fig. 5A**). From our images, we then calculated the total spheroid area (**Fig. 5B**), spheroid ferret diameter (**Fig. 5C**), and spheroid circularity (**Fig. 5D**). Spheroid circularity was not found to be statistically significantly different across groups (*p* > 0.01); however, both spheroid area (*p* = 0.0051) and spheroid Feret diameter (*p* = 0.0002) were statistically significantly different across Blst but not weight percent. Overall, we observed a decreasing trend with area and Feret diameter as Blst increased.

**Figure 5.**
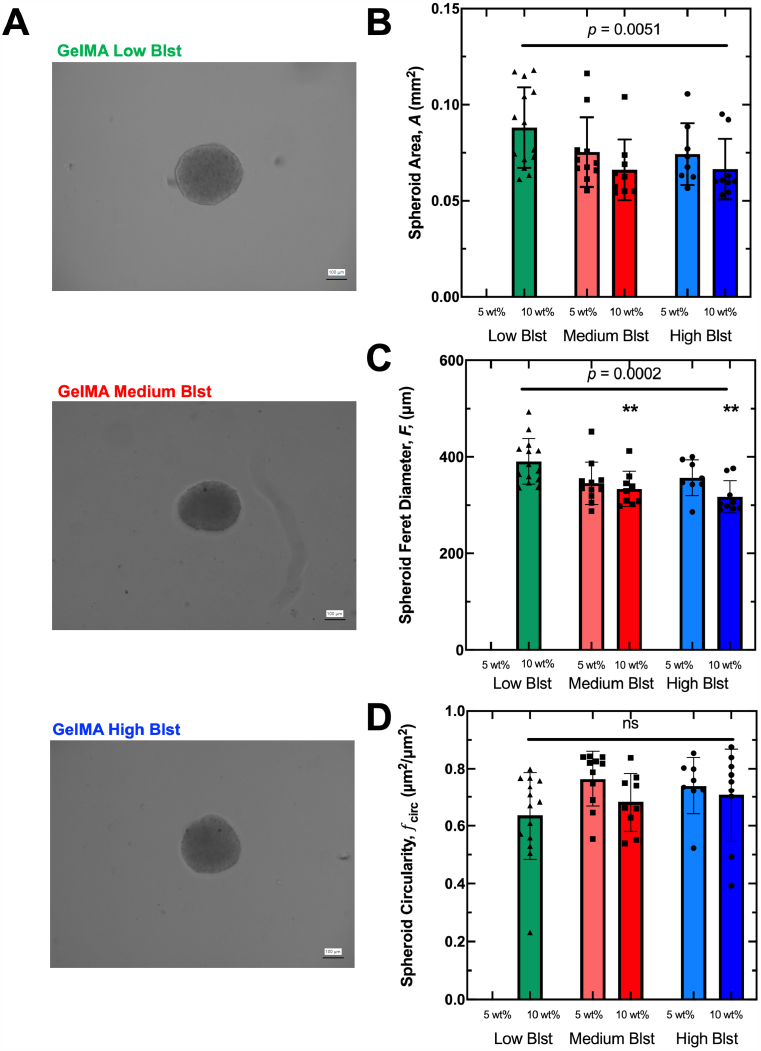
Effect of gelatin bloom strength on Swan71 trophoblast spheroid morphology in gelatin methacryloyl (GelMA) hydrogels. **A**. Representative phase contrast images of encapsulated Swan71 trophoblast spheroids (4,000 cells/spheroid) in 10 wt% GelMA hydrogel variants. Scale: 100 *μ* m. **B**. Average spheroid area, **C**. Spheroid Feret diameter, and **D**. Spheroid circularity immediately after spheroid encapsulation (Day 0). Data presented as mean ± standard deviation (n = 9-14 spheroids per condition). Data analyzed using an ordinary main effects only two-way ANOVA and Tukey posthoc with significance set to p<0.01. Significance of two-way ANOVA depicted as horizontal bar. P-values describe variation from bloom strength (Blst). **: different from low blst 10 wt%, p < 0.01. ns: not significant.

We next plotted spheroid and mechanical properties versus Blst to determine the relationship between Blst and these features. We identified clear relationships between Blst and hydrogel mechanical properties and demonstrated that Blst is related to trophoblast spheroid area and Feret diameter by an exponential decay function (**Fig. 6A**). Previous studies demonstrated that for bulk gelatin gels, Blst shows a linear relationship with elastic modulus^8^. Interestingly, for GelMA, we observed a lognormal trend with a best fit line of R^2^ = 1.0 between Blst and elastic modulus and viscoelastic ratio (**Fig. 6B**).

**Figure 6.**
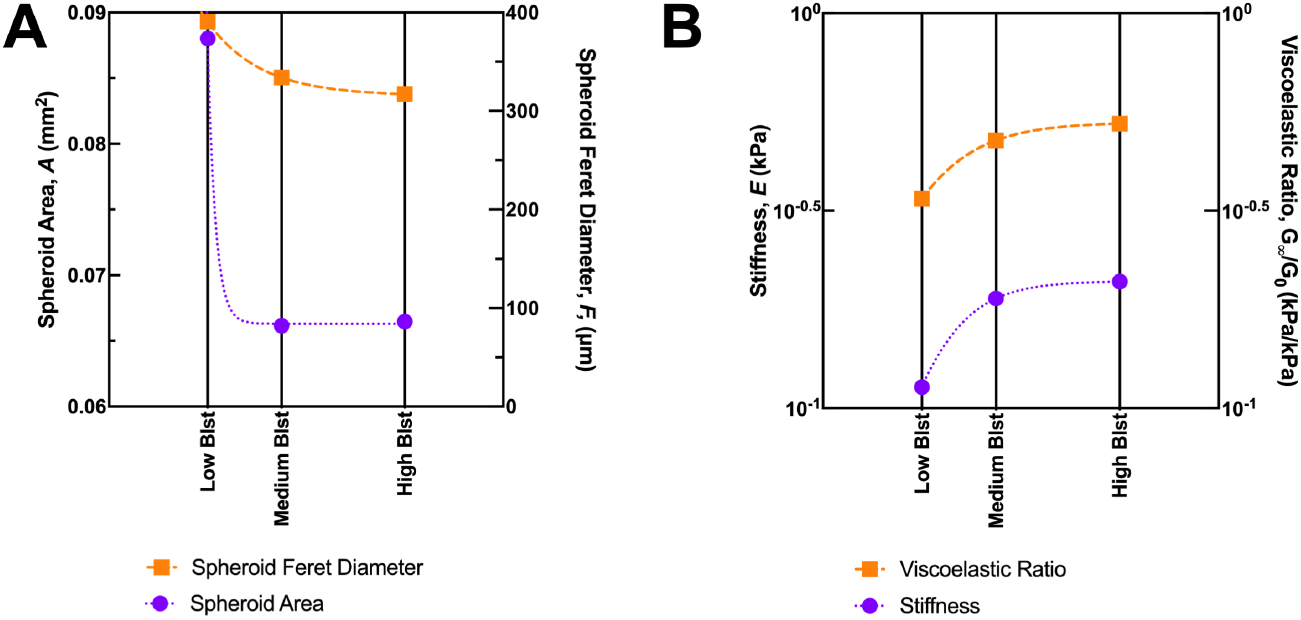
Best fit lines to describe **A**. the relationship between bloom strength (Blst) and spheroid area and Feret diameter (exponential decay) and **B**. stiffness and viscoelastic ratio (lognormal) for 10 wt% GelMA hydrogels of varying Blst.

## 4. Discussion

In this work, we define the role of Blst on GelMA mechanical properties and describe the role of these properties in the context of trophoblast spheroid encapsulation. We synthesized three GelMA variants of low, medium, and high bloom strength. To assess the effects of only Blst, we kept the degree of functionalization, photoinitiator concentration, and UV crosslinking time the same while only changing Blst and weight percent. We systematically quantified hydrogel mechanical properties through spherical nanoindentation and then encapsulated trophoblast spheroids into these hydrogel variants and quantified spheroid area, Feret diameter, and circularity. We identified clear relationships between Blst and hydrogel mechanical properties and Blst and trophoblast spheroid area and Feret diameter.

From the nanoindentation experimentation, we determined stiffness, viscoelastic ratio, diffusivity, permeability, and pore size of our hydrogel variants. We determined that both Blst and weight percent influenced both stiffness and viscoelastic ratio; however, only Blst affected diffusivity, permeability, and pore size. Hydrogel stiffness ranged between approximately 150 – 200 Pa, consistent with previous studies^18,19^. Stiffness increases with Blst and increases with weight percent for hydrogels of the same Blst. Viscoelastic ratio, the ratio between the equilibrium and instantaneous moduli, ranges from 0 to 1 and portrays the degree of viscoelasticity of a material, where *G*_∞_ ⁄*G*_0_ = 0 represents a viscous fluid and *G*_∞_ ⁄*G*_0_ = 1 represents an elastic solid. The viscoelastic ratio of the hydrogel variants fell between 0.37 and 0.53, indicating that GelMA hydrogels are viscoelastic at the microscale. Viscoelastic ratio increased with Blst and increased with weight percent for hydrogels of the same Blst. Diffusivity, permeability, and pore size were also determined from nanoindentation. Diffusivity is a time-dependent material property calculated from permeability and stiffness. Diffusivity ranged from 10^-12^ to 10^-11^ m^2^/s which is consistent with GelMA hydrogels^5^. Intrinsic permeability describes the transport of solvent molecules through the material and is affected by the pore geometry of the material. Permeability of the hydrogels ranged from 10^-17^ to 10^-16^ m^2^ and we determined pore size from permeability and determined the pore size ranged from approximately 4 nm (Medium and High Blst) to 10.9 nm (Low Blst) which generally falls in the range of previous studies^22–24^.However, a limitation of our study is that pore sizes larger than the indenter radius cannot be measured using nanoindentation.

We then determined hydrogel crosslinking density from mass swelling ratio calculations and demonstrated decreasing crosslinking density with increasing polymer weight percent, consistent with previous studies^18^. We determined that the mass swelling ratio was between approximately 11% to 15% for the 10 wt% variants and 25% to 27% for the 5 wt% variants which is within the range of previous mass swelling ratios for GelMA hydrogels^3^. We then determined that weight percent, and not Blst, demonstrated a significant effect on hydrogel crosslinking density.

Higher Blst is correlated with the triple-helical content and the compressive modulus of gelatin^6,8^. Our nanoindentation studies agree with previous studies in that GelMA stiffness increased with Blst. Blst also correlates with gelatin purity and this seemed to also correlate with GelMA purity. The Low Blst GelMA variant was more variable compared to the Medium and High Blst, when assessing the nanoindentation data. Furthermore, the 5 wt% Low Blst variant did not polymerize even with increased UV polymerization time, unlike the Medium and High Blst 5 wt% hydrogels. The Low Blst hydrogel variant had the lowest stiffness and viscoelastic ratio and the highest diffusivity, permeability, and pore size compared to the Medium and High Blst hydrogels. Because of these properties, we then observed the largest spheroid area, Feret diameter, and circularity for the Low Blst condition compared to the Medium and High Blst conditions. These data suggest that softer hydrogels or hydrogels with higher pore size allow for greater spheroid spreading, at least in the context of trophoblast spheroids. This may perhaps be mitigated through a softer, more porous matrix which allows the spheroid to occupy more space within the hydrogel compared to a stiffer matrix with smaller pores. Although, a limitation of our data is that we cannot decouple the effects of Blst and specific hydrogel properties without additional experimentation. Ongoing studies include systematic hydrogel fabrication varying a variety of hydrogel properties (Blst, degree of functionalization, UV intensity, weight percent) to decouple these properties and allow for more comprehensive studies on GelMA hydrogel effects and cellular behavior.

## 5. Conclusions

GelMA is a highly versatile and tunable material for tissue engineering and biomaterials research. Here, we utilize a systematic approach to assess the role of Blst on GelMA hydrogel properties. Blst affected all hydrogel properties studied, including stiffness, viscoelastic ratio, diffusivity, permeability, and pore size, except crosslinking density which was instead affected only by polymer weight percent. Interesting, Blst affected the morphology of trophoblast spheroids at encapsulation, which suggests that the material properties may influence cellular behavior immediately after encapsulation in the material. In identifying clear relationships between bloom strength, hydrogel mechanical properties, and trophoblast spheroid morphology, we demonstrate that bloom strength should considered when designing tissue engineered constructs.

## Acknowledgements

The authors would like to thank Dr. Huafang Li for assistance with ESEM training in the IMSE Core Facility at Washington University in St. Louis. This research was supported in part by the National Institute of Diabetes and Digestive and Kidney Diseases of the National Institutes of Health under award number 1T32DK120497-01A1 (SGZ).

We describe contributions to the manuscript using the Contributor Roles Taxonomy (CRediT)^14^:

*Conceptualization-SGZ, MLO*

*Methodology-SGZ*

*Formal Analysis-SGZ (Nanoindentation; ATR-FTIR; All Statistics), EF (NMR), HFR (Trophoblast Data), DMF (Nanoindentation)*

*Investigation-SGZ (GelMA Synthesis; Hydrogel Synthesis; ATR-FTIR; Trophoblast Experiments), EF (NMR), SSK (Hydrogel Synthesis; Nanoindentation; Mass Swelling), DMF (Nanoindentation)*

*Resources-MLO*

*Data Curation-SGZ*

*Writing: Original Draft-SGZ*

*Writing: Review and Editing-SGZ, EF, SSK, DMF, HFR, SPZ, MLO*

*Visualization-SGZ*

*Supervision-MLO*

*Project Administration-MLO*

*Funding Acquisition-MLO, SGZ*

