## Supplemental Information for "Evaluation of gelatin bloom strength on gelatin methacryloyl hydrogel properties"

### Supplemental Information:
### Systematic evaluation of gelatin bloom strength on gelatin methacryloyl hydrogel properties

**Samantha G. Zambuto[1,2,3], Samyuktha S. Kolluru[3,4], Eya Ferchichi[5], Hannah F. Rudewick[6], Daniella M. Fodera[7], Kristin M. Myers[8], Silviya P. Zustiak[5], Michelle L. Oyen[2,3]**

[1] Department of Obstetrics and Gynecology
Washington University School of Medicine, St. Louis, MO 63130

[2] Department of Biomedical Engineering
[3] Center for Women's Health Engineering
Washington University in St. Louis, St. Louis, MO 63130

[4] The Institute of Materials Science & Engineering
Washington University in St. Louis, St. Louis, MO 63130

[5] Department of Biomedical Engineering
Saint Louis University, St. Louis, MO 63103

[6] Department of Chemical Engineering
Texas A&M University, College Station, TX 77843

[7] Department of Biomedical Engineering
Columbia University, New York, NY

[8] Department of Mechanical Engineering
Columbia University, New York, NY

**Corresponding Author:**
Michelle L. Oyen
Dept. of Biomedical Engineering
Center for Women's Health Engineering
Washington University in St. Louis
1 Brookings Drive
St. Louis, MO 63130


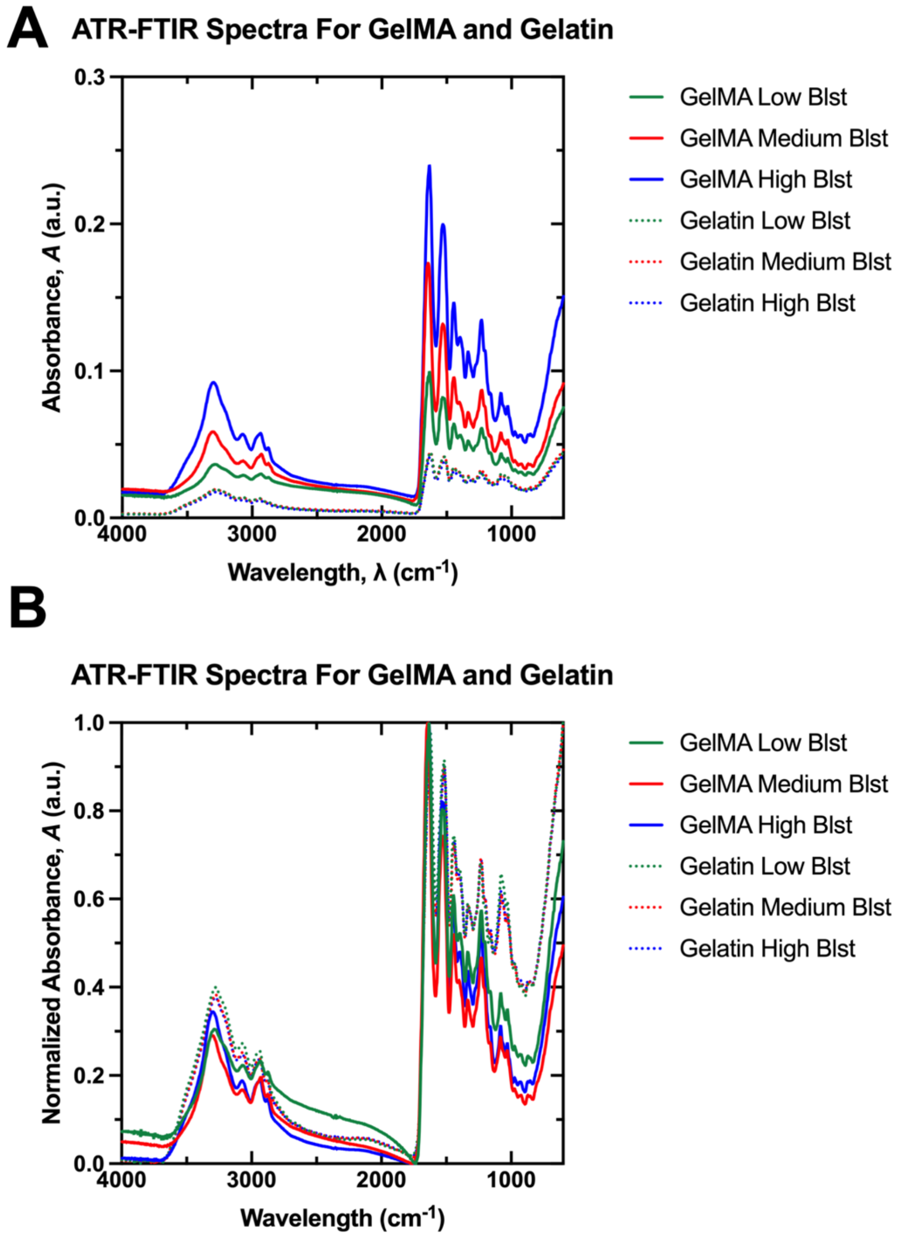


**Supplemental Figure 1. A.** Raw (4000 – 600 $cm^{-1}$) and **B.** normalized ATR-FTIR spectra for gelatin and GelMA variants.

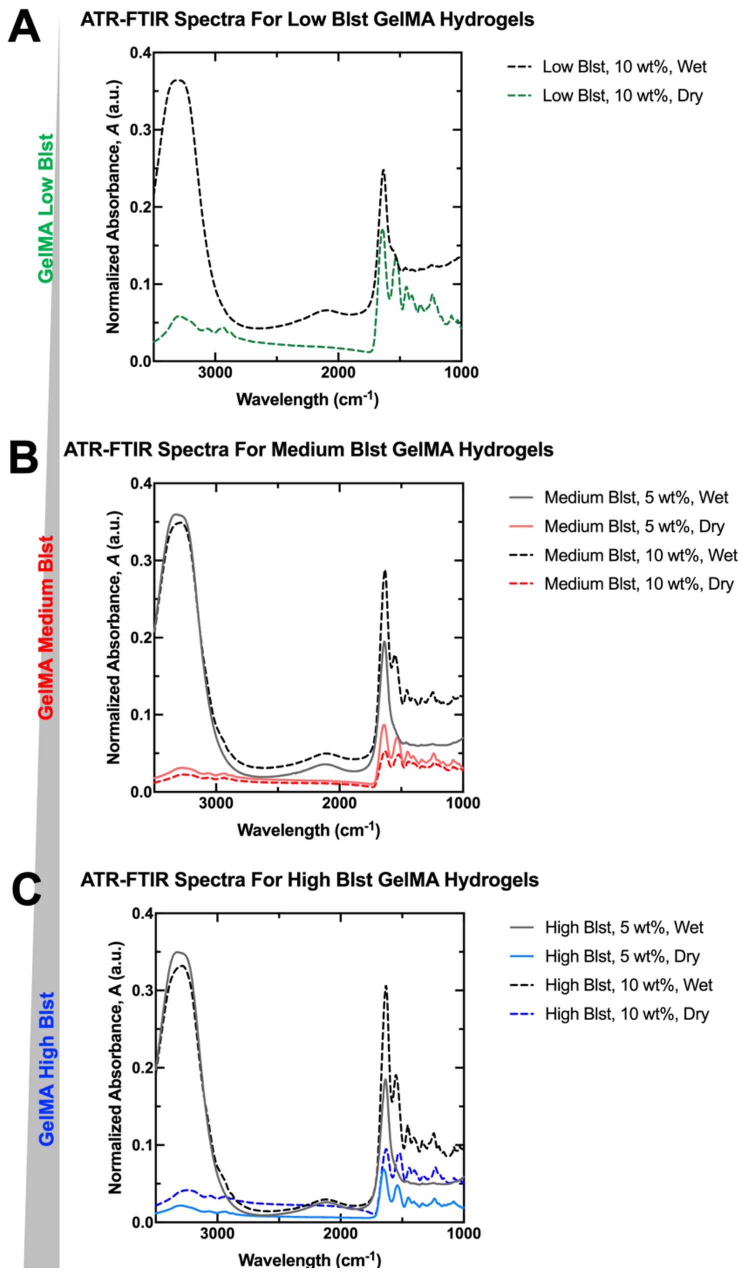


**Supplemental Figure 2.** Normalized ATR-FTIR of hydrated (wet) and lyophilized (dry) crosslinked gelatin methacryloyl hydrogels for **A.** low bloom strength (Blst), **B.** medium bloom strength, and **C.** high bloom strength for 5 wt% and 10 wt%. Low bloom strength, 5 wt% did not polymerize so no data is shown.

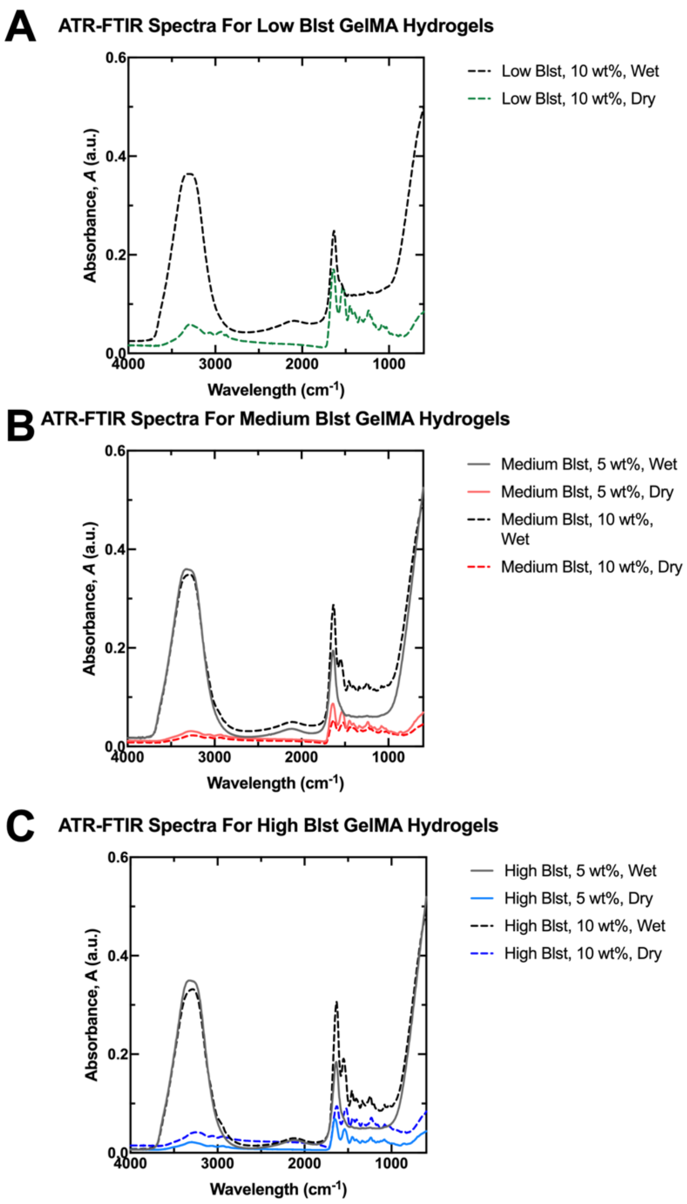


**Supplemental Figure 3**. Raw ATR-FTIR of hydrated (wet) and lyophilized (dry) crosslinked gelatin methacryloyl hydrogels for **A.** low bloom strength (Blst), **B.** medium bloom strength, and **C.** high bloom strength for 5 wt% and 10 wt%. Low bloom strength, 5 wt% did not polymerize so no data is shown.

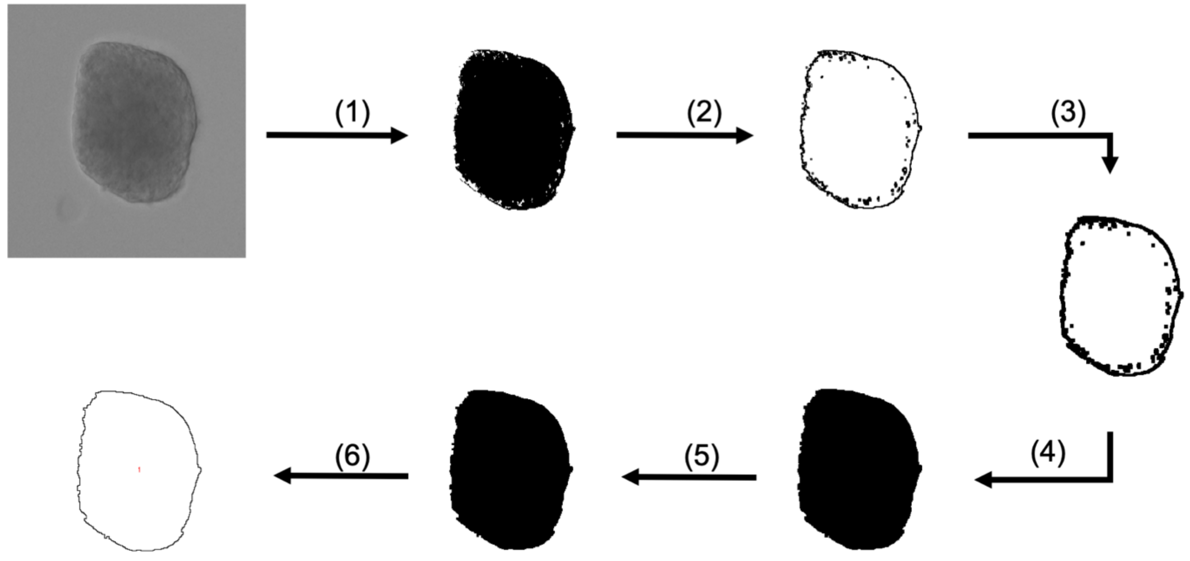


**Supplemental Figure 4.** FIJI macro pipeline for trophoblast spheroid analysis. (1) Binarization (2) Find Edges (3) Dilation (4) Fill holes (5) Erosion (6) Analyze particles (Final measurement yielding area, feret diameter, and circularity).
